## Supplemental Data 1 for "Structural and dynamic changes in P-Rex1 upon activation by PIP_3_ and inhibition by IP_4_"

### Ribbon Map of P-Rex1 (% deuteration)

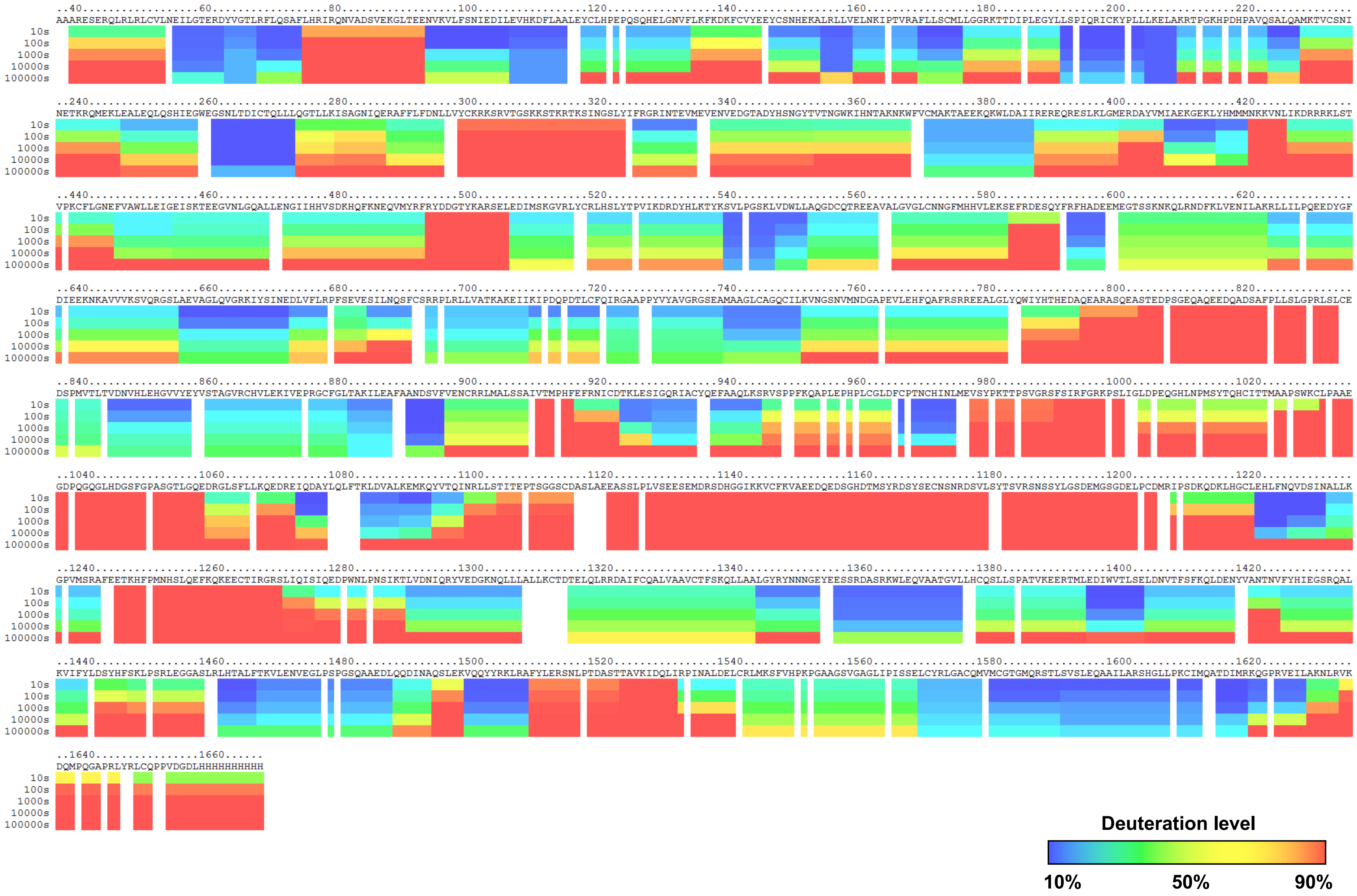

### Ribbon Map of P-Rex1 in P-Rex1•$IP_4$ Complex (% deuteration)

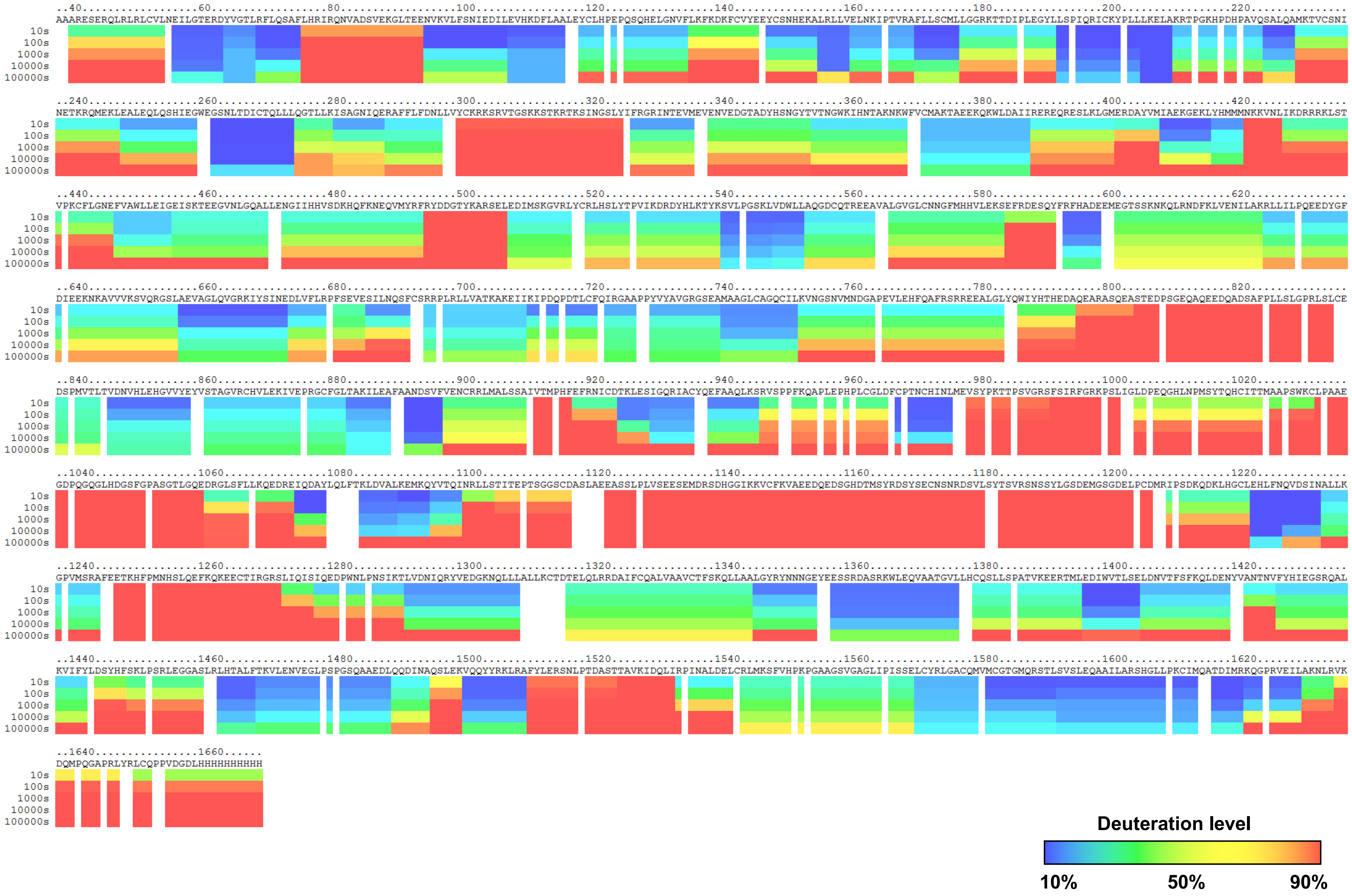

### Influence of $IP_4$ on Exchange in P-Rex1 (% deuteration)

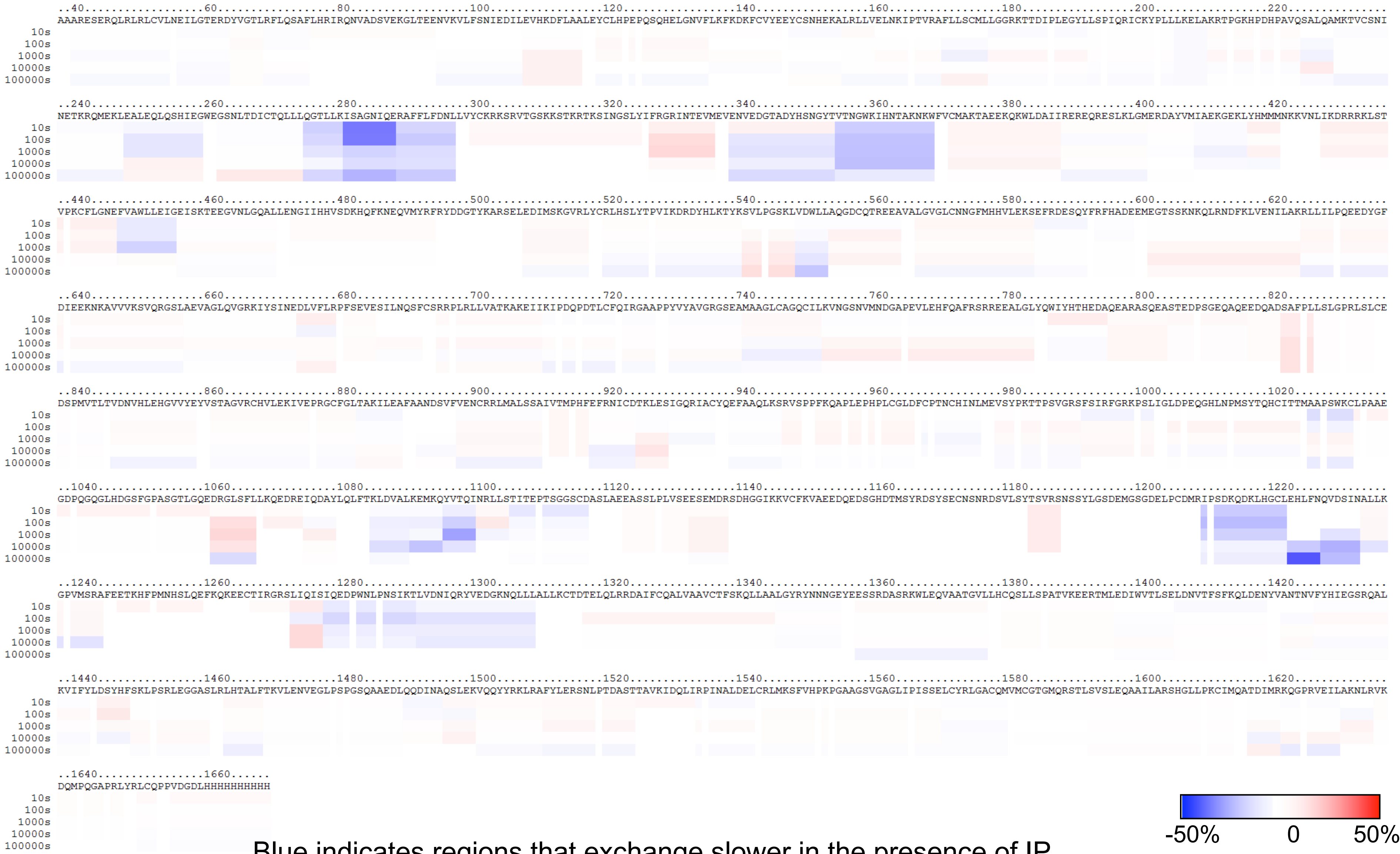


Blue indicates regions that exchange slower in the presence of $IP_4$.
Red indicates regions that exchange faster.
