## Supplemental Data 2 for "Structural and dynamic changes in P-Rex1 upon activation by PIP_3_ and inhibition by IP_4_"

### Ribbon Map of P-Rex1 (% deuteration)

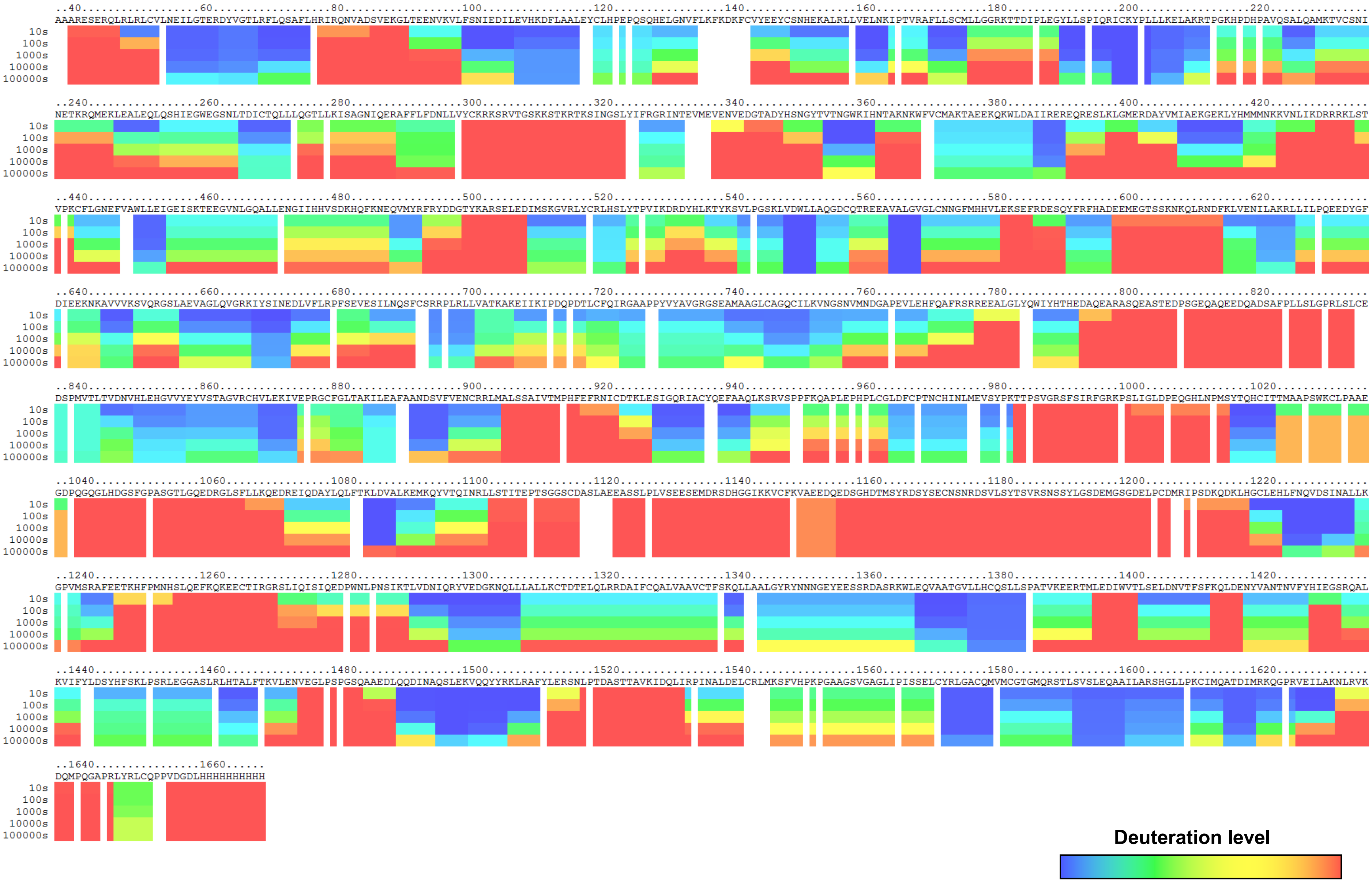

### Ribbon Map of P-Rex1 in the presence of $PIP_3$-containing liposomes (% deuteration)

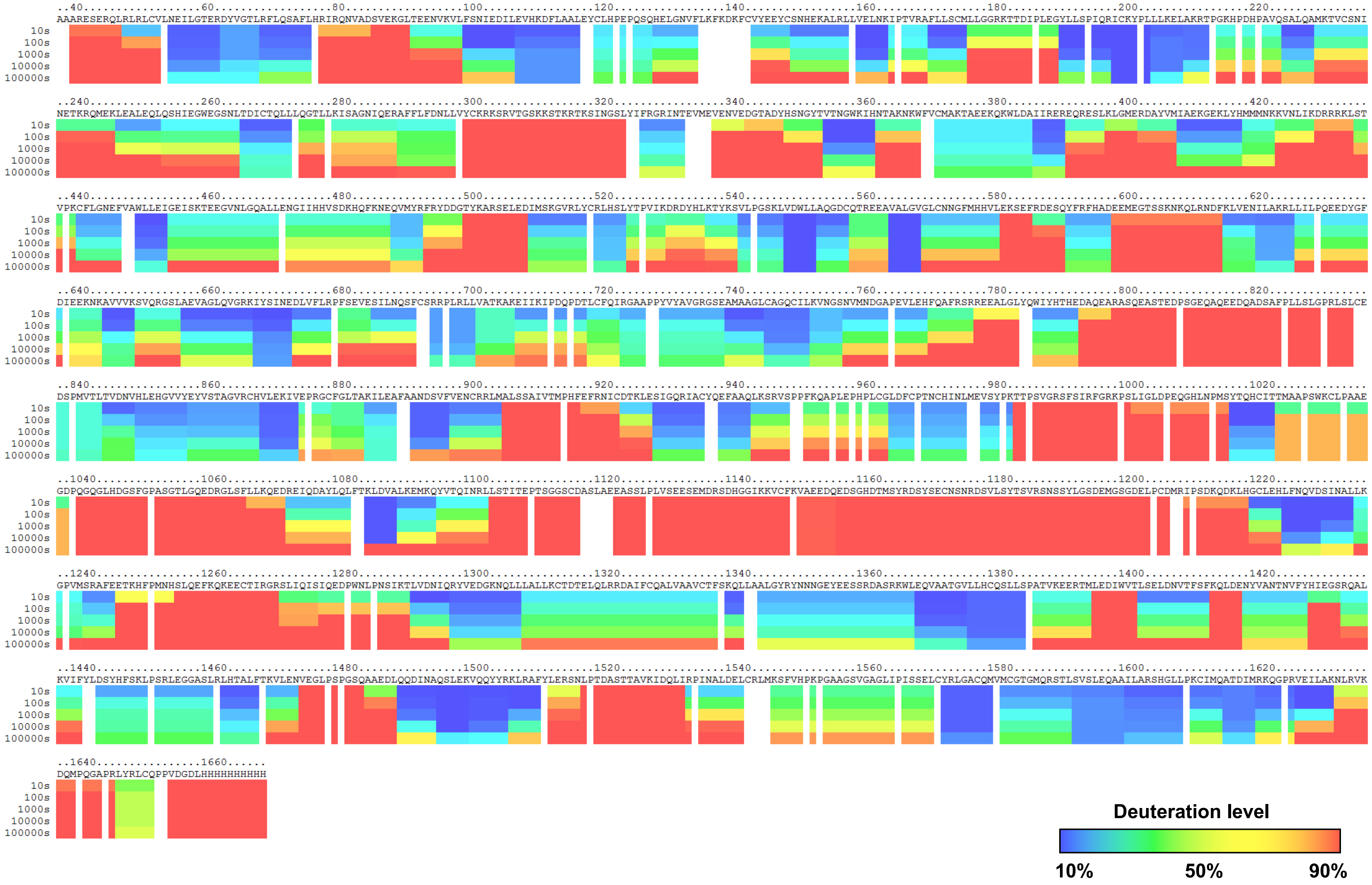

### Ribbon Map of P-Rex1 in the presence of liposomes without $PIP_3$ (% deuteration)

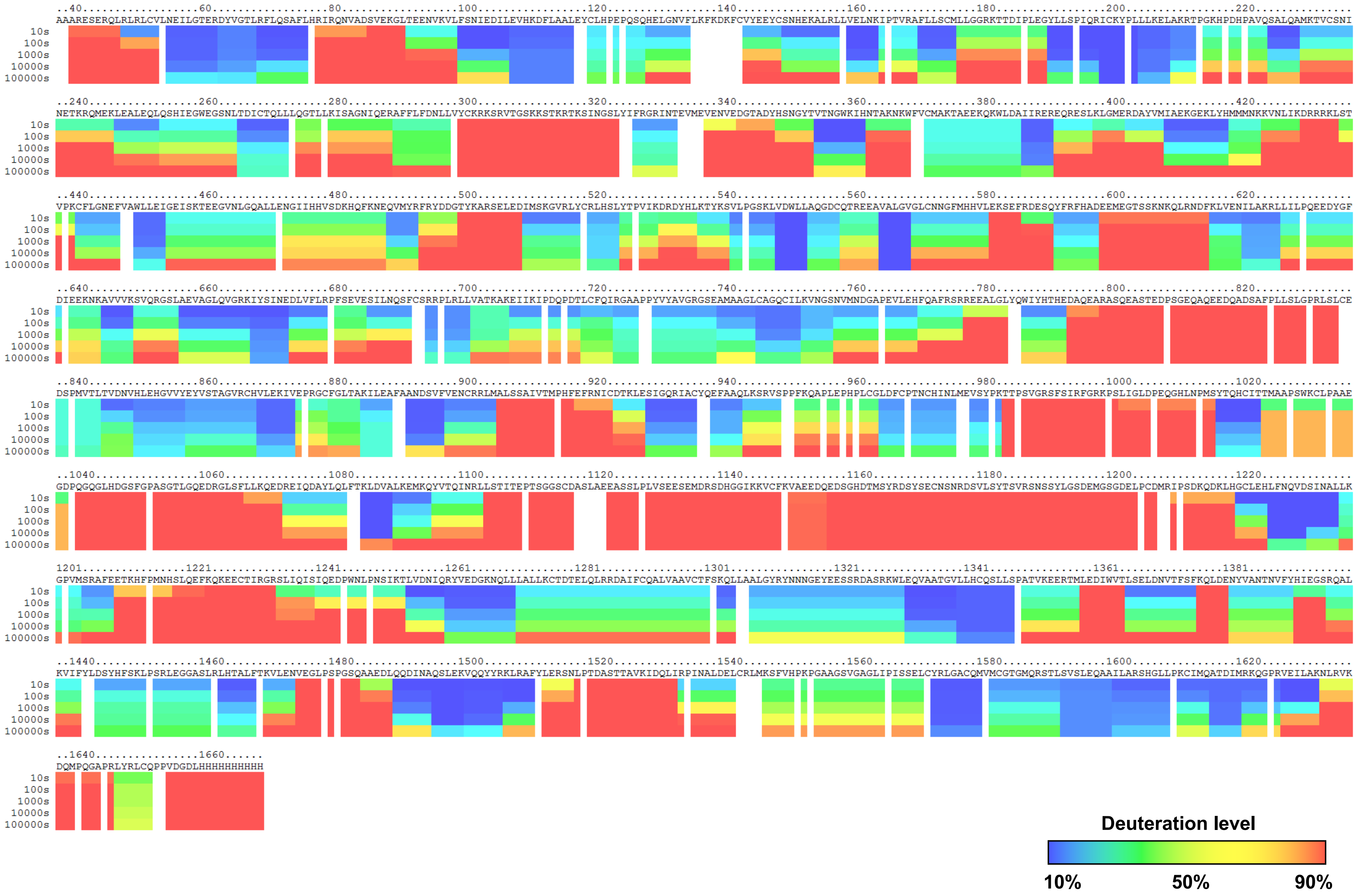

### Difference Map: P-Rex1–$PIP_3$-containing liposomes Minus P-Rex1 (% deuteration)

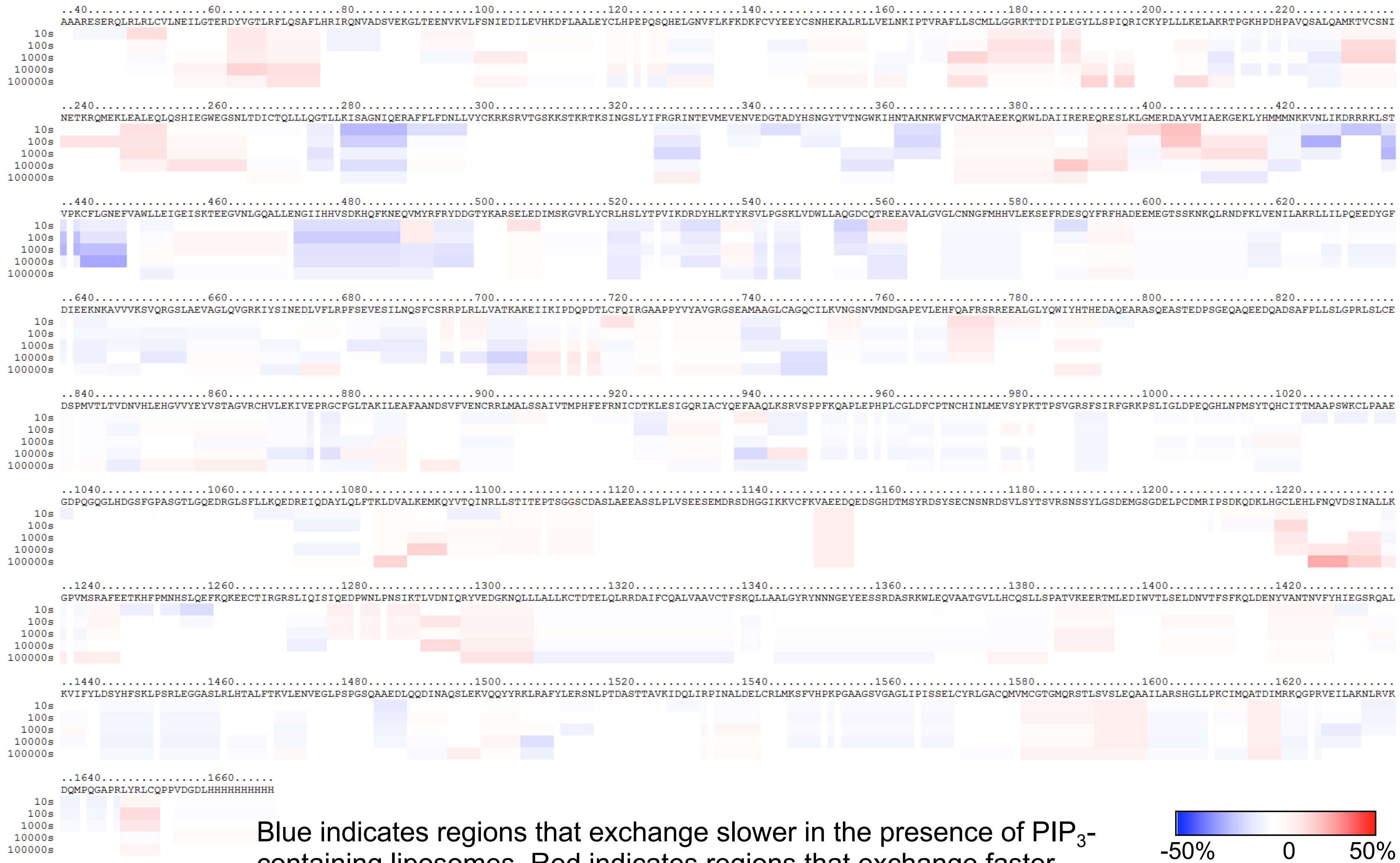


Blue indicates regions that exchange slower in the presence of $PIP_3$-containing liposomes. Red indicates regions that exchange faster.

### Difference Map: P-Rex1–liposomes Minus P-Rex1 (% deuteration)

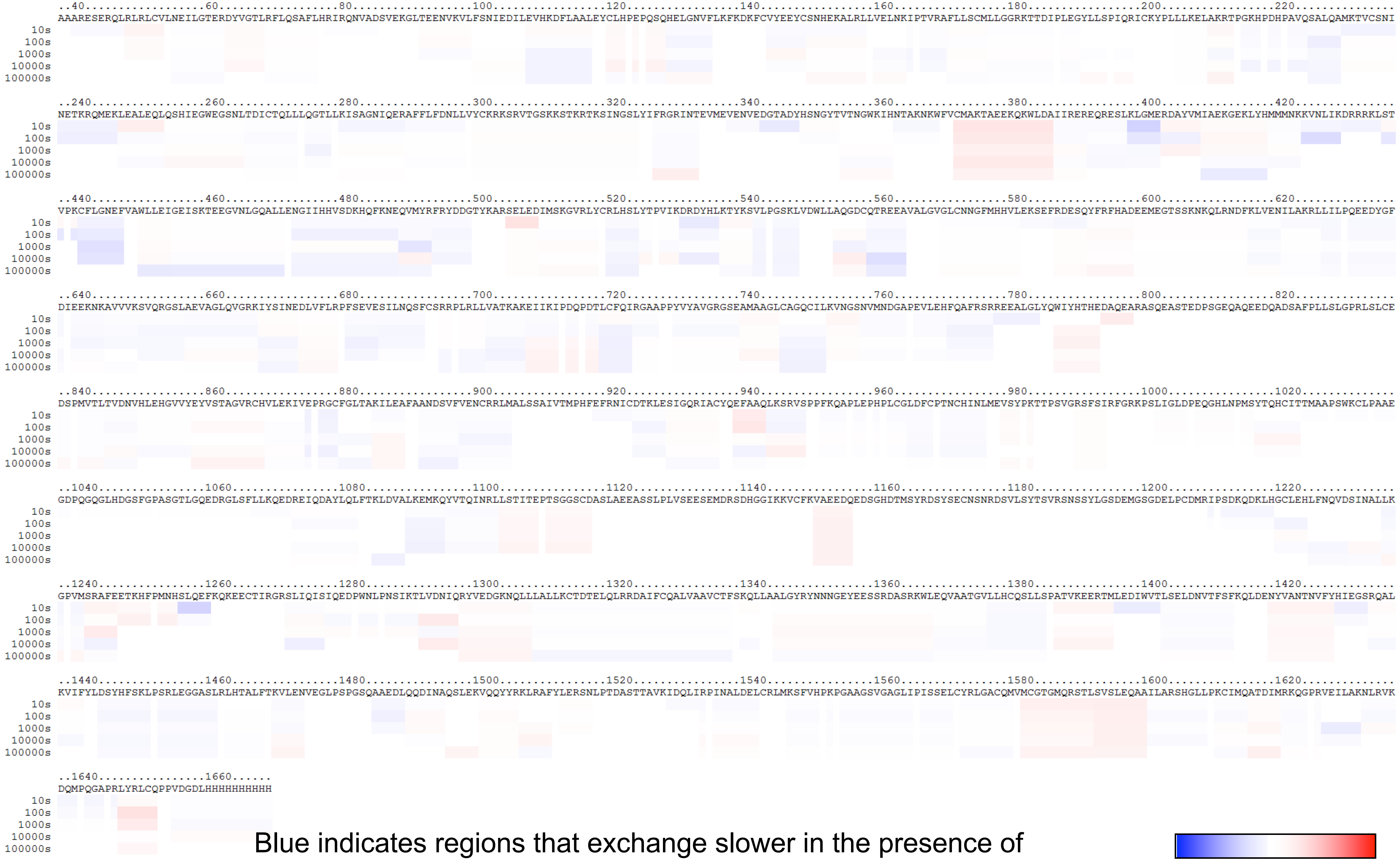


Blue indicates regions that exchange slower in the presence of liposomes without $PIP_3$. Red indicates regions that exchange faster.

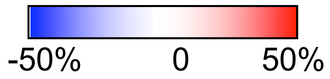
